# Quantifying spinal cord vascular permeability in the mouse using intravital imaging

**DOI:** 10.1101/2021.06.09.447701

**Authors:** M.E. Da Vitoria Lobo, David O Bates, Kenton P Arkill, R.P. Hulse

## Abstract

Sensory perception and motor dexterity is coordinated by in part distinct anatomical centres in the spinal cord. Importantly the spinal cord is the first modulatory relay hub for coordinating sensory and motor inputs to allow control of an organisms response to a sensory experience and to orientate proprioceptive outputs. This is whilst communicating with higher centres within the brain to undertake greater complex neurophysiological function such as pain perception. This begins to outline the complexity of the nervous system communication. To allow this integral system to function efficiently neuronal homeostasis needs to be maintained with energy expenditure matched by proficient delivery of nutrients. This factor introduces the vascular system that extensively interacts in a multifaceted manner with differing aspects of the nervous system. Part of this multi-factoral interaction is through the heterogenic cellular makeup of the vascular network that delivers and modulates the molecular transport of such nutrients to spinal cord tissues, but also controlling penetration and migration of harmful pathogens and agents. Therefore the spinal cord is susceptible to any alterations in the microvessel integrity (e.g. vascular leakage) and/or function (e.g. cessated blood flow) of this vascular network, which principally occurs in times of pathology. Typically investigations into microvessel function have utilised histological and/or tracer based *in-vivo* assays. Methodologies such as evans blue extravasation have been used inconjunction with *in-vitro* cell biology assays such as transwell assays to determine microvessel integrity or function that only provides snapshots of developing vasculopathy. Adopting *in-vivo* imaging approaches, allow for real time functional measurements of the ongoing physiological function within the spinal cord, providing direct measurement of the vascular processes in play, including vascular architecture, blood flow and/or permeability. This technique in mouse allow for direct visualisation of cellular and/or mechanistic influence upon vascular function through utilising disease, transgenic and/or viral approaches. This combination of attributes allows for in depth real time understanding of the function of the vascular network within the spinal cord.

## 1. Introduction

The nervous system consists of a wide degree of differing cellular systems, which work harmoniously to allow for control of integral physiological systems. The vascular system is one part of this, playing a crucial role in the context of neurophysiology and pathology. The vascular system has a multitude of differing roles through providing essential provision for neuronal survival and function through delivering nutrients to the neural tissues through blood perfusion of large arterioles through to small capillaries. Furthermore, in addition it also plays a pivotal role in protecting the nervous tissue from potential damage by preventing any toxic agents or pathogens from invading the tissue. There is an extensive cellular diversity within the vascular network in the nervous system, with cell types consisting of endothelial cells, pericyte, and astrocytes. The interplay between these cell types are essential in providing support to the neural tissues that are integral to an organisms survival [1–3]. Increasingly the health of this vascular system is becoming apparent to be involved with the pathogenesis of neurodegenerative disease, with damage to the vasculature strongly correlating with neuronal damage such as in multiple sclerosis[1][2], alzheimers [3, 4] and stroke[5, 6]. In these instances numerous techniques and models have been adopted, utilising human and rodents tissues for the investigation of this neurovascular system.

In relation to the spinal cord, it is essential to the processing of incoming sensory and motor information enabling the organism to interact with its surrounding environment. Disturbances in microvessel function are prominently associated, though not exclusively, with exacerbated vascular leakage in the spinal cord in neuropathological circumstances resulting in impaired motor coordination and sensory perception. Commonly, a breakdown in the endothelial junctional barrier is a causative factor in the development of neurodegenerative disease (e.g. multiple sclerosis [7]) and motor impairment (e.g. spinal cord traumatic injury[8]) as well as disturbances in nociceptive processing and resulting chronic pain manifestation (such as in arthritis, traumatic injury [9, 10]). Typically these studies have investigated a breakdown in the integrity in the endothelium that subsequently leads to increased levels of vascular permeability [9] and tissue penetrance of invading cell types[11]. This arises due to decreases in endothelial cell production of junction proteins and/or secretion of enzymatic proteases from infiltrating and/or resident cell types that can breakdown these junctional proteins enabling cell infiltration [12–14].

Current approaches to investigate the integrity and functionality of the blood spinal cord barrier have focussed upon primarily histological evaluation of spinal cord tissue from rodents, humans and transgenic/viral models [2, 15]. Utilising fluorescent based antibody immunohistochemistry as well as fluorescent reporter cellular labelling via transgenic mouse and/or viral approaches allows for the appreciation of the microvessel architecture and function within nervous as outlined. Furthermore, measurement of vascular leakage is also determined through immunofluorescent based assays following infusion of fluorescently labelled agent that can be identified in tissues through histological tissue preparation or alternative via quantifying the amount of the solute that penetrates the tissue. Quantification of vascular permeability following infusion of a fluorescent tracer (e.g. tritc-dextran, Evans blue[9]) or fluorescently labelled cell types (e.g. monocyte[16]) administered either via intravenous or systemic approaches can be evaluated [9, 17]. These techniques provide information regarding how the structure of the microvessels networks are influenced by differing experimental interventions. This allows investigation into fundamental physiological processes and disease pathogenesis, as well as how the microvessel function can also be influenced by such factors. However, these studies solely provide a time stamped snapshot of the functionality and integrity of the microvasculature in the nervous tissues. This results in the process of development and continued progression of this vasculopathy not being fully appreciated, whilst changes to the inherent cellular signalling that impacts greatly upon the physiological process is undefined.

Development of improved optical imaging platforms has forged the development and advancement of intravital imaging of nervous tissues. The advantages of this approach allows for acquisition of an array cellular physiological responses such as vessel leakage, using a variety of experimental tools and sensors (e.g. measure cellular calcium influx in situ), in real time. In reference to the vascular network, vessel architecture can be determined whilst simultaneous acquiring a variety of physiological measures of this system such as vessel perfusion and permeability. Despite the physical difficulties of stabilising and accessing the spinal cord, invivo imaging of the rodent spinal cord has recently been utilised by a number of groups [18–20] to determine microvessel function in health and disease. Here we present the utilisation of invivo spinal cord imaging methodology to quantify in real time vessel permeability via leakage of fluorescent tracers in the spinal cord.

## 2. Materials

### 2.1. Ethical Approval and Animals used

All experiments involving animal were performed in accordance with the Unitell anid Kingdom animals (Scientific procedures) Act 1986 and local ethical review board. Animals had *ad libitum* access to standard chow and housed under 12:12h light:dark conditions. All studies for immunohistochemistry and intravital imaging studies were performed in 12 adult male C57.bl6 mice (25-30g; Charles River).

### 2.2. Immunohistochemistry

Adult male mouse terminally anaesthetised via intraperitoneal injection of Ketamine (50mg/kg) and Medetomidine hydrochloride (0.5mg/kg), Wheat germ agglutinin AlexaFluor conjugated 555 (WGA555, 4mg/kg, 100uL in sterile PBS per mouse, Thermoscientific) was intravenously injected via the tail vein. 15 minutes post injection of WGA555, cardiac perfusion fixation was performed using 4% paraformaldehyde. The lumbar spinal cord was extracted and submerged in 4% paraformaldehyde overnight at 4°C. This was followed cryoprotection by submersion in 30% sucrose overnight at 4°C. Tissue was cryosection at 50m thickness and mounted on superfrost slides or stored at -80 °C until required for processing. Processed spinal cord slides were incubated with DAPI (made up in water) for 5 minutes in a hydration chamber at room temperature and a coverslip was mounted with vectoshield (Vector labs). Process slides were imaged on a Leica SP8 confocal microscope.

### 2.3. Anaesthesia Induction and Laminectomy

Adult male mice were terminally anaesthetised via intraperitoneal injection of Ketamine (50mg/kg) and Medetomidine hydrochloride (0.5mg/kg). Anaesthestia was regularly monitored for anaesthetic depth (e.g. foot and corneal reflex), with anaesthesia maintained via continued intraperitoneal injection of Ketamine (50mg/kg) and Medetomidine hydrochloride (0.5mg/kg) where appropriate. Following anaesthesia induction mouse body temperature (∼37oC) was maintained through feedback control system using a heat pad and a rectal probe. Access to the spinal cord was performed via a laminectomy of the thoracic/lumbar region of the spinal cord. The dorsal surface of the vertebral column was shaved, all hair removed and the region cleaned/sterilised with chlorohexidine. An insertion along the vertebral column on the dorsal surface was made using a scalpel blade approximately 2.0cm in length. Skin was separated from underlying tissues using blunt dissection exposing the muscle and connective tissue. Muscle and connective tissue were removed to expose the vertebrae and the vertebral processes. Vertebral processes were removed including transverse processes along with any remaining muscle and connective tissue from the lateral aspects of the vertebrae. A laminectomy was performed between T12-L1 to expose the lumbar segments L3-L5 of the spinal cord. Following exposure of the spinal cord, the tissue was kept hydrated with sterile saline.

### 2.4. Intravital Spinal Cord Imaging Preparation

An inhouse window chamber was assembled and attached to mouse spinal vertebra as outlined. The window chamber consists two sets of side pillars and attachment bars that clamp either side of the vertebra. This provides a fixed configuration to enable attachment to the microscope securely allowing for stable image capture. The face plate for the window chamber, which possesses a recessed insert for a glass coverslip, is a attached to the side pillars, holding the window chamber in position. A silicone elastomer (Kwik Sil, World Precision Instruments) is used to fill the gap in the face plate to immerse the spinal cord. A glass coverslip (diameter = 5mm, thickness = #0, Warner Instruments) is inserted into the recess in the face plate of the window chamber and pressed against the silicone elastomer. A bubble fee seal was ensured and left to allow the elastomer to cure If bubbles were present or coverslip not secure, the coverslip was removed and elastomer was reapplied with coverslip inserted.

### 2.5. Intravital Imaging Fluorescent Vessel Labelling

To identify the endothelium in the spinal cord, AlexaFluor 555 conjugated wheat germ agglutinin, (WGA555, 4mg/kg, 100uL in sterile PBS per mouse, Thermoscientific) was intravenously injected via tail vein. The mouse was left for 15 min and positioned on the microscope stage and the spinal cord was imaged using a confocal microscope (Leica SPE). Vessels were identified as those that contained WGA555 labelled endothelium. Following identification of labelled vessels, permeability was measured using sodium fluorescein. Sodium fluorescein recommended 100mg/ml in sterile PBS, Sigma-Aldrich was inject via intraperitoneal injection. Immediately after sodium fluorescein injection, the spinal cord was imaged (minimum of 120s with a frame rate of 10 frames/s). All settings regarding confocal laser configuration (e.g. gain and offset) were maintained for all experiments. Furthermore, velocity of blood flow was determined by imaging sodium fluorescein filled vessels to track blood cell movement via shadows (areas of no fluorescent molecule in the lumen)

### 2.6. Statistical Analysis

All image processing was performed using Fiji (https://imagej.net/Fiji)[21], whilst GraphPad Prism v8 was used for data analysis. All data is presented as mean ±standard error of mean. Acquired images were captured (.LIF) and exported as using Leica LAS software (acquired image precision of either 256 x 256 pixels or 512 x 512 pixels). These were converted into .TIFF file format. Using Fiji identified vessels radius (identified using WGA555 staining) was determined using calibrated freehand tool. Sodium fluorescein vessel leakage was measured using mean fluorescence intensity in a chosen region of interest (ROI) that spanned across the vessel wall; including fluorescein vessel intensity inside the vessel lumen and adjacent tissue outside the vessel wall. Whilst in comparison the mean fluorescence intensity in the adjacent tissue outside the vessel follows a linear increase over time. This allows the initial sodium fluorescein permeability 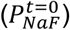 to be calculated by EQ. 1 and illustrated in fig 4A[22]:

**Figure 1.**
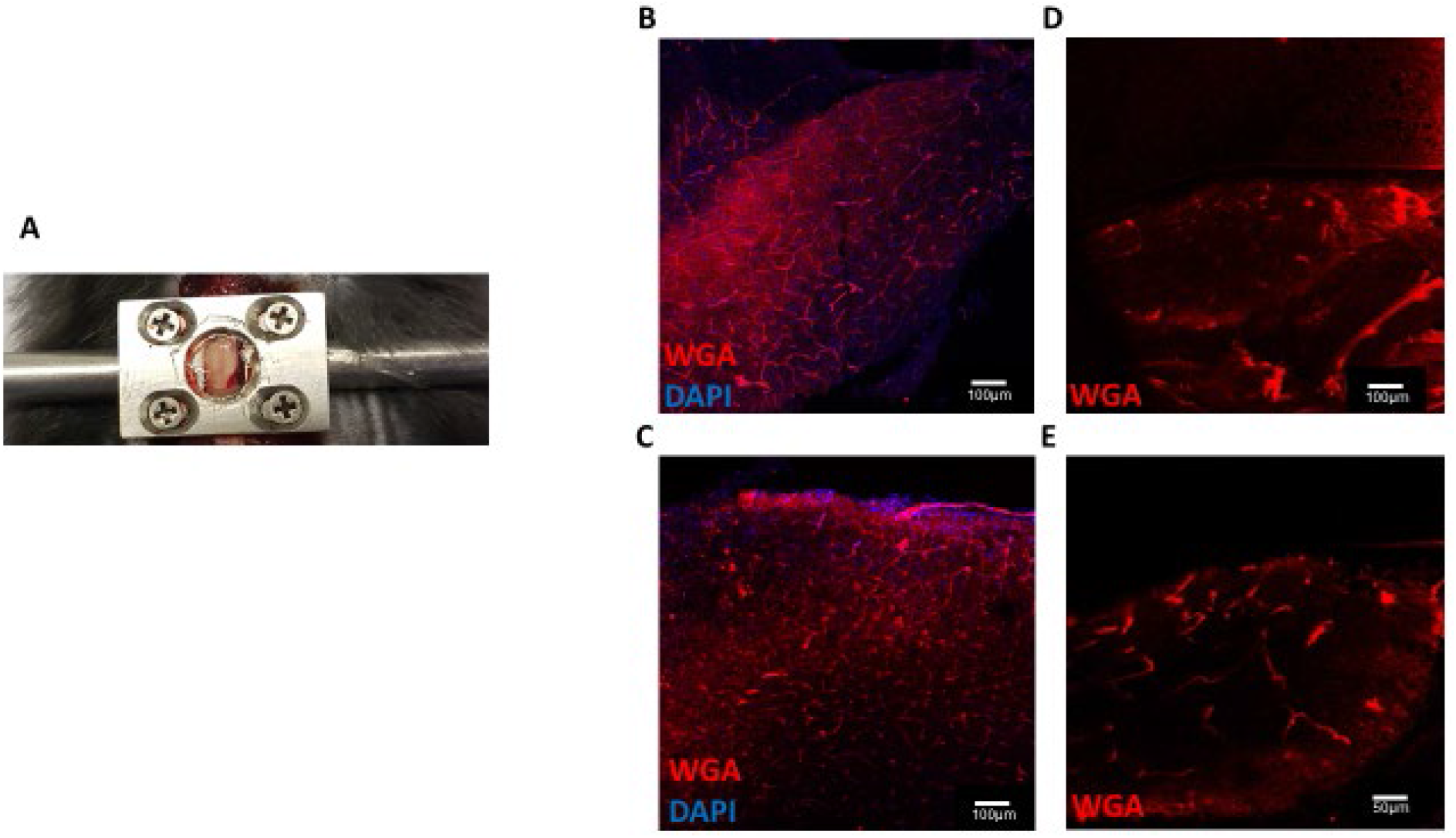
Spinal cord microvessel imaging invivo and fluorescentl labelling of the endothelium in the spinal cord [A] Representative image of the window chamber in situ plus glass coverslip and silicone elastomer in place. Cryosectioned lumbar [B] brain and [C] spinal cord sections (50µm) prepared from C57.bl6 adult mice injected with Wheat Germ Agglutin Alexafluor 555 (4mg/kg), delivered via intravenous tail vein injection allow visualisation of the endothelium in the central nervous system (scale bar = 100µm). Wheat Germ Agglutin Alexafluor 555 (4mg/kg) administered via intravenous tail vein injection [D & E] allows visualisation of microvessels in the spinal cord invivo via intravital imaging.

**Figure 2.**
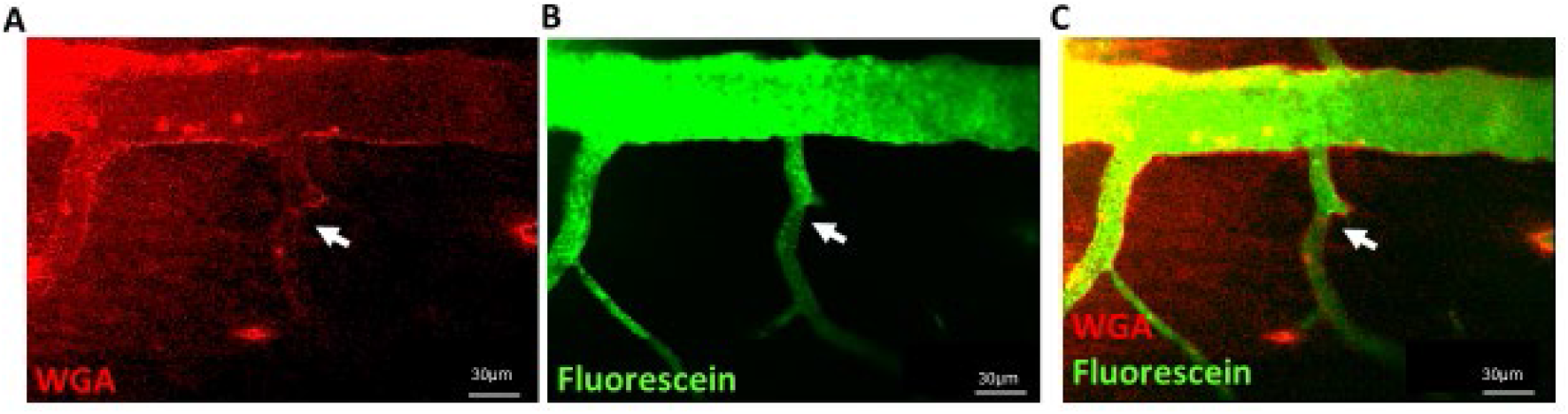
Fluorescently labelling of the endothelium in the spinal cord Intravenous injection via the tail vein of Wheat Germ Agglutin Alexafluor 555 (4mg/kg) [A] allows visualisation of microvessels in the spinal cord invivo via intravital imaging. To measure vessel permeability sodium fluorescein is injected via intraperitoneal injection. Representative example of a [B] WGA 555 labelled (Red) vessels (arrows) in the spinal cord and [C] sodium fluorescein perfused (Green; Overlay provided in [D]).

**Figure 3.**
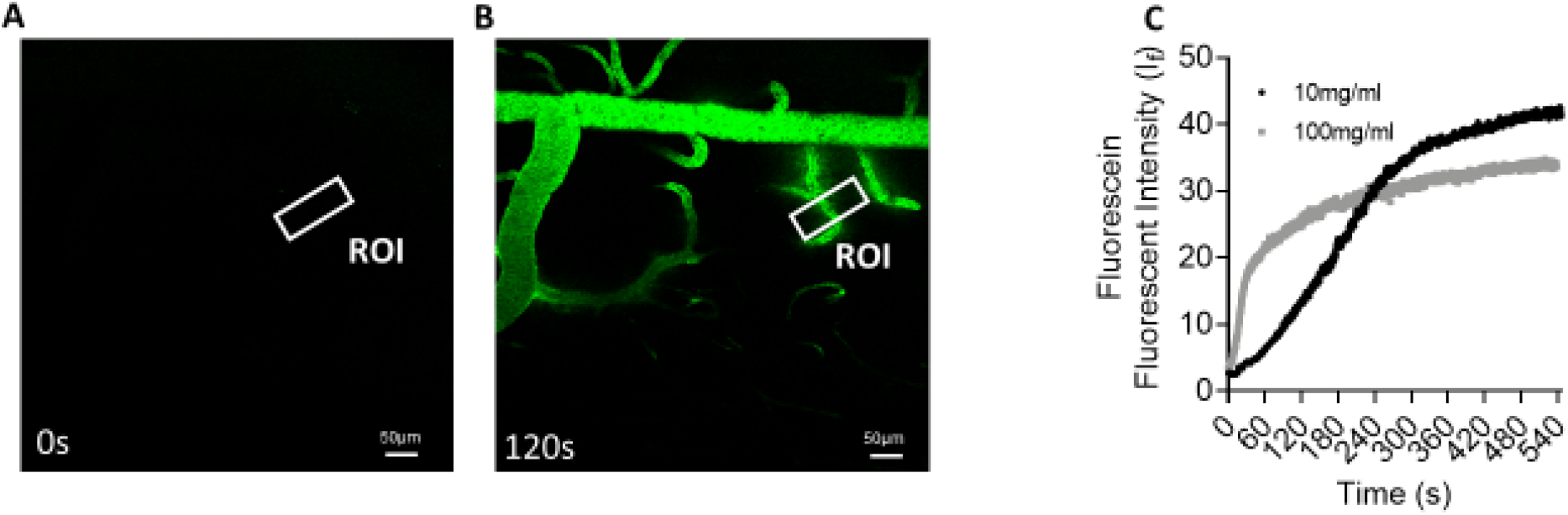
Sodium fluorescein perfused microvessels in the spinal cord Sodium fluorescein is administered via intraperitoneal injection. Representative examples following intraperitoneal injection of 100mg/ml sodium fluorescein at Time [A] 0s and [B] 120s demonstrating vessels are perfused with sodium fluorescein by 120s. [C] Mean fluorescence intensity in the region of interest (ROI) over time for 10mg/ml and 100mg/ml Sodium fluorescein demonstrate 100mg/ml provides a steeper rate of vessel perfusion.

**Figure 4.**
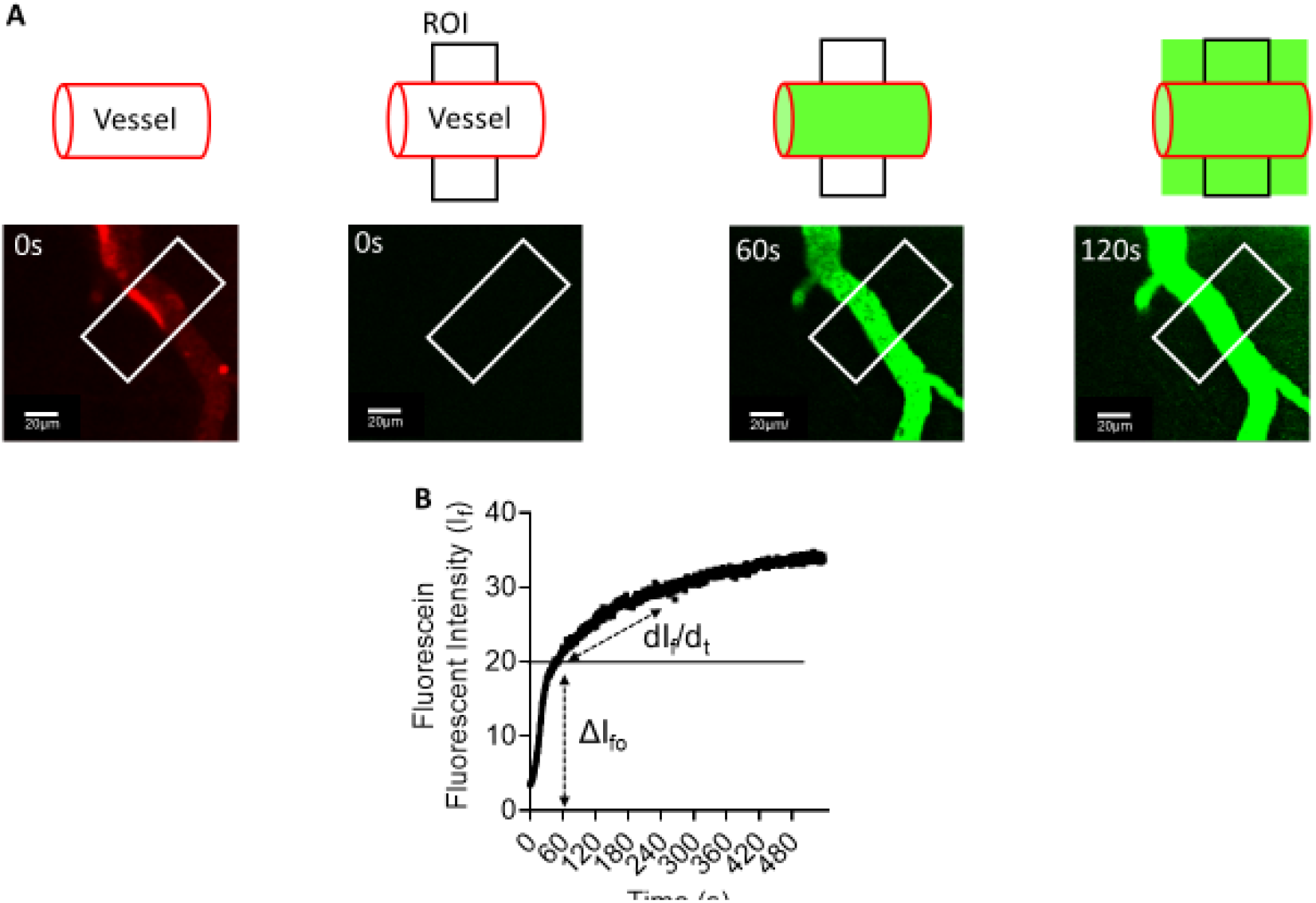
Sodium fluorescein permeability quantification [A] A region of interest (ROI) is identified across an identified vessel labelled with WGA555 (scale bar = 200µm). Sodium fluorescein is injected via intraperitoneal injection and fills the vessel lumen over 120s. [B] Mean fluorescence intensity is displayed over time demonstrating a steep rate of increase mean fluorescence intensity in the region of interest (ROI), with initial mean fluorescence intensity change (ΔI_f_) depicting change from T=0s to mean fluorescence intensity plateau (approximately 60s). Subsequent rate of increase in mean fluorescence intensity represents vessel leakage of sodium fluorescein over time (dI_f_/d_t_).

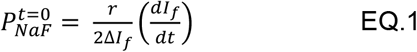

Where

*I*_*f*_ = mean fluorescence intensity of ROI

*r* = radius of vessel

*ΔI*_*f*_ *=* Mean fluorescence intensity from the luminal vessel fluorescein bolus

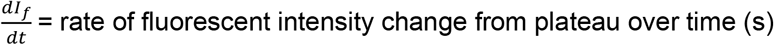.

It is not possible to determine the capillary hydrostatic pressure directly; however, the velocity can be determined using kymographs of cell trafficking through the vessel of known radius. The velocity can be related to volumetric flow (*F*) as EQ2.

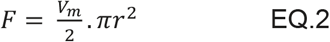

Where *V*_*m*_ *i*s the maximum velocity. Whilst full quantification is impossible, particularly with blood, *F c*an be approximated by:

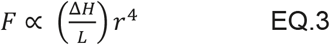

Where Δ*H i*s the change in Hydrostatic pressure along the length (*L*) of vessel, i.e. 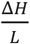 is the hydrostatic pressure at the ROI. Therefore, if desired, particularly for changes in pathology if the proportionality of *V*_*m*_ to *r*^*2*^ *i*s constant then any permeability changes are likely to be predominately structural changes rather than an upsteam mechanistic effect such as vasoconstriction.

## Results

A real time measure of vascular permeability in the spinal cord was determined using fluorescent labelling of the spinal cord microvessels captured via a spinal cord window chamber in an anesthetised mouse (Fig.1A). A laminectomy was performed to expose the lumbar spinal cord and a custom-made window chamber (consisting of a face plate that houses the coverslip and vertebral side clamps) was secured to the vertebral column. To enable identification of the vasculature in the spinal cord, fluorescent labelling was utilised to stain the endothelium. Wheat germ agglutinin binds to sialic acid residues present in the glycocalyx layer covering the endothelium as observed in-vivo previously [23]. In spinal cord cryosections processed from mice that were administered with WGA555 (4mg/kg) via tail vein injection, the blood vessel endothelium was identified with WGA55 positive labelling in the dorsal horn of the spinal cord (Fig. 1B & C). In anaesthetised mice prior to evaluation of vascular permeability, WGA555 endothelial staining in the spinal cord was used to identify vessels in the spinal cord (Fig.1D & E). Following identification of the microvessel with WGA555 labelling (Fig. 2A), vessel permeability in the spinal cord was determined. Intraperitoneal injection of sodium fluorescein was used to monitor vessel permeability from the vessels in the spinal cord (Fig.2B, representative merge example Fig.2C) allowing monitoring of increased distribution and accumulation in the lumen of the vessel as well as vascular leakage. Post injection sodium fluorescein distribution in the vessel network of the spinal cord was acquired from 0s (point of injection; Fig.3A) over 2 minutes (Fig.3B), with sodium fluorescein administration delivered initially at either one of two doses (10mg/ml or 100mg/ml). In all consequent follow up studies 100mg/ml was administered.

In an identified WGA555 labelled vessel (Fig. 4A), solute permeability was determined via sodium fluorescein injection. Fluorescent intensity of sodium fluorescein was measured by recording the increased mean fluorescence intensity in the ROI against time (Fig. 4A & B). The measure of solute permeability is the determination of a fluorescently labelled molecule to move across a region of vessel wall (as outlined via the white box in Fig. 4A) ie the quantity of solute moving across the wall over time. This rate of solute movement follows the assumption that the number of fluorescently labelled molecules spread either side of the vessel wall at T=0s are equal and therefore the linear increase in fluorescence intensity in the tissue mileu (120s dI_f_/d_t_) is determined from the initial step to fill the vessel lumen (ie at 60s, ΔI_fo_). This is presented as initially following sodium fluorescein administration, there is a rapid increase in fluorescence intensity within the vessel lumen, which plateaus and stabilises (Fig. 5A, ΔI_fo_). However, fluorescent intensity outside the vessel wall in the adjacent tissue increases in a linear manner (Fig. 5A, dI_f_/d_t_), indicative of solute transport across the vessel wall. To note, this measure of solute permeability is dependent upon movement of fluorescently labelled molecules to transport into the surrounding vessel milieu, with the assumption of undisturbed hydrostatic pressure. Hydrostatic pressure was assumed to be constant as there were no fluctuations in red blood vessel velocity (Fig. 5B) or identified vessel diameter across a known length of capillary (Fig. 5C).

**Figure 5.**
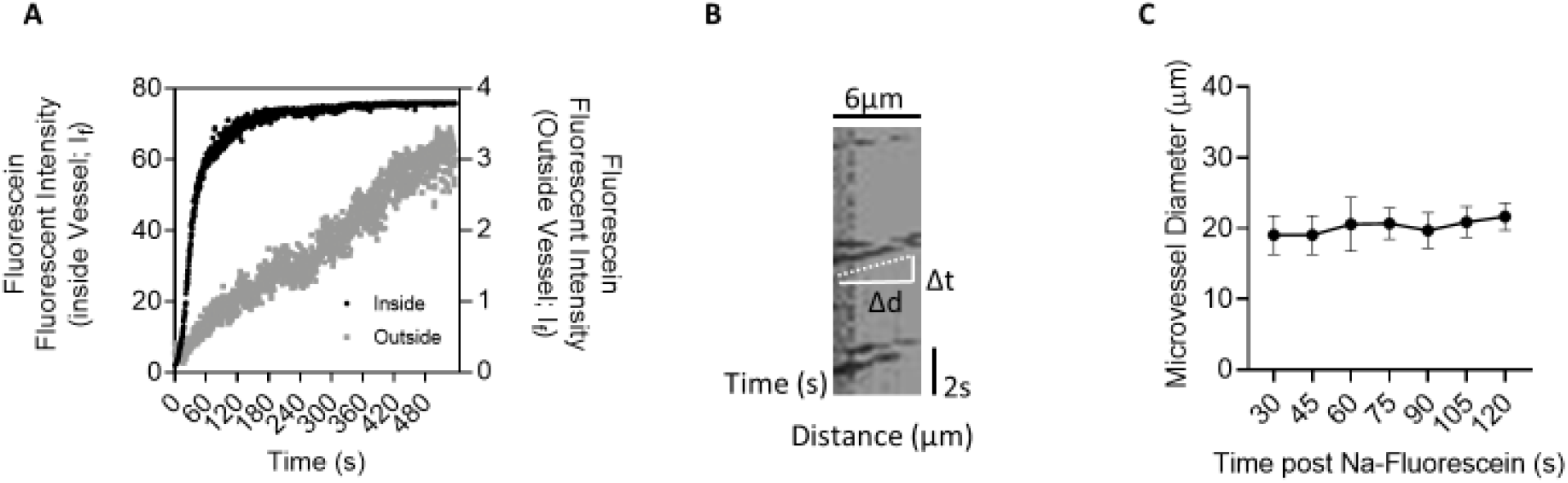
Vessel permeability and quantification of blood flow. [A] Represents the mean fluorescence intensity of sodium fluorescein in the vessel lumen (black) over time depicting steep rate of increase until plateau accompanied by a linear increase in mean fluorescence intensity in the adjacent tissue outside the vessel wall (grey). [B] Kymograph depicting unaltered trajectory of blood cell movement through a vessel highlighting no change in vessel diameter or red blood cell velocity and [C] no change in vessel diameter.

## Discussion

The spinal cord is a complex cellular system, which is a crucial regulator of somatosensory and proprioceptive processing, fundamental physiological systems for an organisms function and survival. Unfortunately in a number of differing pathological states the vascular network that feeds the spinal cord is susceptible to damage from an array of differing diseases, which is displayed as alterations in vessel network architecture and/or function. In many instances of neurological disease such as with multiple sclerosis [1] and inflammatory pain [10, 15], alterations in this vascular network is underpinned by elevated neuroinflammatory processes. This is typically displayed as increased infiltration of inflammatory cell types (e.g. macrophage infiltration) and/or elevated vascular leakiness (e.g. increased penetration of IgG or fluorescent tracers), physiological processes that are underpinned by molecular alterations of the cellular composition and architectural changes in the microvessels within the spinal cord [10–12, 15]. Experimental approaches to date typically rely upon inherently time locked studies, which only provides a snapshot of the systems in play in relation to vessel function though offering inights into the concept of increased vascular permeability.

Here we present a study that takes advantage of advances in optical imaging to in real time measure vessel function, principally focussing upon vascular permeability. The endothelium is a semi-permeable barrier that coordinates tissue homeostasis through the movement of molecules and cells transported in the blood. As highlighted above, vessel permeability is the physiological process that is determined by net capillary differences across a membrane of solute and water whilst considering the impact of hydrostatic pressure [24, 25]. Furthermore, vessel structural parameters including interjunctional proteins (e.g. Claudin, Occludin, Zonula Occludens 1) and an endothelium luminal carbohydrate rich filter known as the glycocalyx are also factors of consideration when determining molecular insights into vessel permeability [23, 25]. This presented intravital methodology in normal rodent allows for the evaluation of molecular transport across the endothelial wall. Using fluorescent labelling the structural context of the luminal vessel lining is identified, whilst allowing for the tracking of solute movement using a fluorescent labelled molecule to solute permeability. Utilising this approach allows the fundamental process of vessel permeability not to be inferred, but represented by true physiological parameters allowing full appreciation of this biological system in the spinal cord to investigate related aspects of health and disease.

## Acknowledgements

RPH and MD performed the experimental work. RPH, MD, KA and DOB contributed to the conception or design of the work in addition to acquisition, analysis or interpretation of data for the work. RPH, MD, KA and DOB drafted the work or revised it critically for important intellectual content. RPH drafted the manuscript with contributions from all authors. All authors approved the final version of the manuscript, agree to be accountable for all aspects of the work in ensuring that questions related to the accuracy or integrity of any part of the work are appropriately investigated and resolved all persons designated as authors qualify for authorship, and all those who qualify for authorship are listed. We would like to acknowledge Prof Chris Schaffer at Cornell University for advice in relation to design of the window chamber.

